# Autophagy is essential for optimal Fe translocation to seeds in *Arabidopsis*

**DOI:** 10.1101/313254

**Authors:** Mathieu Pottier, Jean Dumont, Céline Masclaux-Daubresse, Sébastien Thomine

## Abstract

Micronutrient deficiencies affect a large part of the world population. They are mostly due to the consumption of grains with insufficient content of Fe or Zn. Both *de novo* uptake by roots and recycling from leaves may provide seeds with nutrients. Autophagy, which is a conserved mechanism for nutrient recycling in eukaryotes, was shown to be involved in nitrogen remobilization to seeds. Here, we have investigated the role of this mechanism in micronutrient translocation to seeds. We found that several *Arabidopsis thaliana* plants impaired in autophagy display defects in nutrient remobilization to seeds. In *atg5-1* mutant, which is completely defective in autophagy, the efficiency of Fe translocation from vegetative organs to seeds was severely decreased even when Fe was provided during seed formation. Combining *atg5-1* with *sid2* mutation that counteracts premature senescence associated to autophagy deficiency and using ^57^Fe pulse labeling, we could propose a two-step mechanism in which iron taken up *de novo* during seed formation is first accumulated in vegetative organs and subsequently remobilized to seeds. Finally, we showed that translocations of zinc and manganese to seeds are also dependent on autophagy. Fine tuning autophagy during seed formation opens therefore new possibilities to improve micronutrient remobilization to seeds.

## Introduction

Metal micronutrients are essential to all forms of life. For instance, iron (Fe) plays a major role in oxido-reduction reactions allowing respiration in mitochondria and photosynthesis in chloroplasts (Nouet *et al.*, 2011). Worldwide, about 2 billion people suffer from Fe deficiency which affects mostly children and women in developing countries (WHO, 2016). Staple food crops are poor sources of Fe and the major place they take in human diet is one of the leading cause for Fe deficiency (Murgia *et al.*, 2012). During the last decade, different biofortification strategies such as fertilization, conventional breeding, and genetic engineering, were pursued in an attempt to increase key micronutrient levels in seeds of crop species (Murgia *et al.*, 2012).

The improvement of micronutrient loading in seeds requires knowledge of micronutrient trafficking pathways within the plant, including uptake from soil and remobilization from senescing organs. To date, most studies on micronutrients have focused on their uptake in root cells, intracellular partitioning, and long-distance transport. Several proteins participate in the mandatory solubilization and the reduction of Fe to Fe^2+^ prior to its uptake by a plasma-membrane specific transporter, AtITR1, in the epidermis of Arabidopsis roots (Robinson *et al.*, 1999; Vert *et al.*, 2002; Santi and Schmidt, 2009; Fourcroy *et al.*, 2014; Lefèvre *et al.*, 2018). Fe is transported to the shoot upon loading in the xylem (Durrett *et al.*, 2007; Morrissey *et al.*, 2009) and translocation to sink tissues is mediated by loading/unloading in the phloem after Fe chelation to nicotianamine (NA) (Schuler *et al.*, 2012). Finally, the oligopeptide transporter 3 (AtOPT3) could transport chelated Fe to ensure Fe loading to seeds (Stacey *et al.*, 2008). During vegetative growth, Fe is mostly directed towards photosynthetic tissues. Fe is crucial for photosynthesis and over 80% of the Fe in mesophyll cells is concentrated in chloroplasts (Shingles *et al.*, 2002). Thus, photosynthetic tissues constitute a large Fe pool potentially available for remobilization during seed filling. However, the mechanisms of micronutrient remobilization from vegetative organs during senescence have received little attention so far (Pottier *et al.*, 2014). Senescence in *Arabidopsis thaliana* is associated to a decrease by 50% of leaf Fe concentration, in parallel with micronutrient filling in seeds (Himelblau and Amasino, 2001). This finding implies that 50% of the micronutrients present in senescent leaves are not remobilized. A better understanding of micronutrient remobilization from senescent organs is therefore likely to highlight new solutions to improve seed micronutrient content. More specifically, mechanisms controlling the availability of nutrients in source organs such as autophagy, could be a matter of great significance for micronutrient loading in seeds (Shi *et al.*, 2012; Pottier *et al.*, 2014).

Autophagy is conserved from yeast to animals and plants. It allows degradation of cytoplasmic components by lysosomal (animals) or vacuolar (plants) internalization mediated by double membrane vesicles called autophagosomes (Reumann *et al.*, 2010). Autophagy promotes energy production, toxic components elimination, and nutrient recycling by discarding aberrant proteins, damaged organelles as well as normal cytoplasmic components when they are no longer useful (Yoshimoto, 2012). Autophagy related genes (*ATG*) involved in autophagosome formation were first identified in yeast (Matsuura *et al.*, 1997). Subsequently, orthologues of *ATG* genes were identified in plants and autophagy deficient plants were characterized (Li and Vierstra, 2012). All autophagy deficient plants exhibit hypersensitivity to carbon and nitrogen starvation pointing to a central role of autophagy for nutrient recycling (Liu and Bassham, 2012; Guiboileau *et al.*, 2013). The up-regulation of *ATG* genes during leaf senescence in *Arabidopsis* suggests a role for autophagy in nutrient recycling at the end of the plant life cycle (Doelling *et al.*, 2002; van der Graaff *et al.*, 2006; Chung *et al.*, 2010; Breeze *et al.*, 2011; Liu and Bassham, 2012). In agreement with this hypothesis, Nitrogen Remobilization Efficiency, i.e. the proportion of nitrogen allocated to seeds through remobilization from vegetative tissues during senescence, is strongly affected in autophagy deficient plants (Guiboileau *et al.*, 2012).

In this work, we investigated the role of autophagy in metal micronutrient resorption from senescent leaves to seeds in *Arabidopsis*. Our results indicate that micronutrient remobilization is impaired in vegetative tissues of autophagy deficient plants. Focusing on ATG5, which is essential for autophagosome formation (Kirisako *et al.*, 2000), we show that autophagy-dependent remobilization is critical for optimal translocation of Fe as well as other metal nutrients from vegetative organs to seeds.

## Material and methods

### Plant material and growth conditions

*Arabidopsis thaliana* (L.) ecotype Columbia-0 (Col-0) [*atg5-1* (SALK_020601), *atg9-2* (SALK_130796), AtATG18a RNAi (RNAi18), *sid2*, *atg5sid2*], and ecotype Wassilewskija (WS) [*atg4* (*atg4a4b-1*), *atg9-1*] have been described previously (Hanaoka *et al.*, 2002; Yoshimoto *et al.*, 2004, 2009; Xiong *et al.*, 2005; Inoue *et al.*, 2006).

The first experiment (experiment 1) including all autophagy deficient plants presented in this study was conducted on sand. Seeds were sawn on sand and plants were watered three times per week for 2 hours with high nitrogen nutritive solution as described in Chardon *et al.* (2010). Plants were grown under the following conditions: 8/16 h, 21/17°C light/dark until 56 days after sowing (DAS) and then transferred to long days (16 h light), maintaining similar day/night temperatures. When plants were dry, seeds were separated from dry remains of vegetative tissues, including leaves, stems, and siliques to determine their respective dry weights and metal contents.

In other experiments (experiments 2 and 3) performed to determine the involvement of autophagy in Fe remobilization, plants were cultivated on a 1/1 mix of sand and perlite, watered three times per week for 2 hours with modified Hoagland solution [0.28 mM KH_2_PO_4_, 1.25 mM KNO_3_, 0.75 mM MgSO4, 1.5 mM Ca(NO_3_), 25 μM H_3_BO_3_, 50 μM KCl, 1 μM ZnSO_4_, 0.1 μM Na_2_MoO_4_, 0.5 μM CuSO_4_, 5 μM MnSO_4_, and 3 mM MES-KOH, pH 5.7]. Ten micromolar Fe was provided as Fe^3+^ chelated to HBED [N,N’-di(2-hydroxybenzyl) ethylene diamine-N,N’-diacetic acid monochloride hydrate; Strem Chemicals, Newburyport, Massachusetts, USA] prepared as described by Lanquar *et al.* (2005). Plants were grown in a climate chamber under the following conditions: 75% relative humidity; 9/15 h, 21/19°C light/dark until 68 days after sowing. Then, plants were transferred to long days (16 h light), maintaining similar conditions. At the onset of flowering (78 DAS) half of plants were maintained in Fe sufficient nutrition (solution containing 10 μM Fe-HBED) and the other half were transferred to Fe deprivation nutrition (solution containing 20 μM of ferrozine) until the end of the plant life cycle. When plants were dry, leaves, seeds, and stems including empty siliques were harvested separately to determine their dry weight and their metal content. In experiment 2, light was enriched in blue and red wavelengths (OSRAM FLUORA, Munich, Germany) and light intensity was maintained to 100 μmol of photon.m^-2^.s^-1^. In experiment 3, which is presented in supplemental figures, white light was provided by TLD 58W 830 and 840 (Philips, Amsterdam, Netherlands) with an intensity of 230 μmol of photon.m^-2^.s^-1^.

### Metal concentration analysis

After weight measurement, dried samples were digested in 2 mL of 70% nitric acid in a DigiBlock ED36 (LabTech, Italy) at 100°C for 1 h, 120°C for 6 h and 80°C for 1 h. After dilution in trace-metal-free water, the metal content of the samples was determined by atomic absorption spectrometry using an AA240FS flame spectrometer or with a MP AES 4200 Atomic Emission spectrometer (Agilent, USA).

### ^57^Fe-labeling

For ^57^Fe labeling, ^57^Fe^3+^ (96.28 Atom %) was prepared and combined with HBED to replace ^56^FeHBED by ^57^FeHBED in the modified Hoagland solution. After four days without watering, plants were placed in presence of ^57^FeHBED modified Hoagland solution for 24 h precisely, at 52 and 54 DAS, during the vegetative phase. After labeling, the pots containing substrate and roots were rinsed four times with ultrapure water and one time with non-labeled solution to discard ^57^Fe. Then, unlabeled modified Hoagland solution was used for the rest of the culture cycle until harvest.

### Determination of ^57^Fe/^56^Fe ratios and remobilization indices

An aliquot of 50 mg of dried and crushed sample was digested in a mixture of nitric acid hydrogen peroxide (ultra-pure reagent grade) on a heated block DigiPrep (SCP Science, Canada) at 90°C during 3 hours and then diluted to 50 mL with ultrapure water. Blank samples were prepared in the same conditions. The measurement of the isotopes ^56^Fe and ^57^Fe contained in these solutions was performed using Inductively Coupled Plasma Mass Spectrometry, ICP MS (Perkin Elmer Nexion, USA). Helium gas was introduced in the dynamic collision cell to prevent from the specific polyatomic interferences caused by argon and calcium oxides. Quality control solutions were prepared from a standard solution containing 100 mg.mL^-1^ of natural sourced iron (Inorganic Ventures) and placed during the whole analytical sequence. The ^57^Fe abundance (A%) was calculated as atom per cent and defined as A% = 100 x ^57^Fe content / (^57^Fe content + ^56^Fe content). The ^57^Fe abundance of unlabeled plant controls (A% _control_) was 0.0241. The ^57^Fe enrichment (E%) of sample was defined as (A% _sample_ – A% _control_). The ^57^FeRSA_sample_ was calculated as (A% _sample_ – A% _control_) / (A% labeling solution – A% control). The 57FeRSAseeds / 57FeRSA(seeds + stems + leaves) ratio was calculated as (E% _seeds_) / [(E% _seeds_ x Fe% _seeds_ x DW _seeds_ + E% _leaves_ x Fe% _leaves_ x DW _leaves_ + E% stems x Fe% stems x DW stems) / (Fe% seeds x DW seeds + Fe% leaves x DW leaves + Fe% stems x DW stems)] where E% is the ^57^Fe enrichment, Fe% is the concentration of Fe in percent, and DW is the dry weight (Gallais *et al.*, 2006; Masclaux-Daubresse and Chardon, 2011).

### Statistical analysis

Experiments were carried out in three to eight independent biological replicates. Data were analyzed with Kruskal-Wallis and Mann-Whitney non-parametric tests for multiple comparisons and pair comparisons, respectively. For multiple comparisons, a Tukey post hoc test was performed when significant differences were detected (*p < 0.05*). Different letters indicate significant differences between samples. All tests were performed using the R software package.

## Results

### Autophagy deficient plants retain higher concentrations of zinc, manganese, and iron in vegetative organs

To investigate the involvement of autophagy in micronutrient remobilization from vegetative organs to seeds during senescence, we first compared metal concentrations in vegetative organs of wild-type and several autophagy deficient plants after the completion of the plant’s life cycle in experiment 1 (Hanaoka *et al.*, 2002; Yoshimoto *et al.*, 2004; Thompson *et al.*, 2005; Xiong *et al.*, 2005; Guiboileau *et al.*, 2012). If remobilization of metal micronutrients to seeds is impaired in plants with compromised autophagy, their concentration in the dry remains of vegetative parts should be increased compared to wild-type. Manganese (Mn), iron (Fe), and zinc (Zn) concentrations were up to 2.5 times higher in dry remains of autophagy deficient plants compared to those of wild-type plants (**Fig. 1**). The highest increases in metal concentrations were consistently observed in *atg5-1* and *atg4a atg4b-1* mutants, which are fully defective in autophagosome formation (Yoshimoto *et al.*, 2004; Guiboileau *et al.*, 2012). In contrast, in dry remains of *atg9.1*, *atg9.2*, and ATG18 RNAi, increases in Mn, Fe, and Zn concentrations were smaller and, in some cases, not significant. This is consistent with the observation that autophagic bodies have been detected in *atg9.1*, *atg9.2,* and ATG18 RNAi plants, indicating that autophagy is not fully compromised in these mutants (Yoshimoto *et al.*, 2004; Guiboileau *et al.*, 2012).

**Figure 1.**
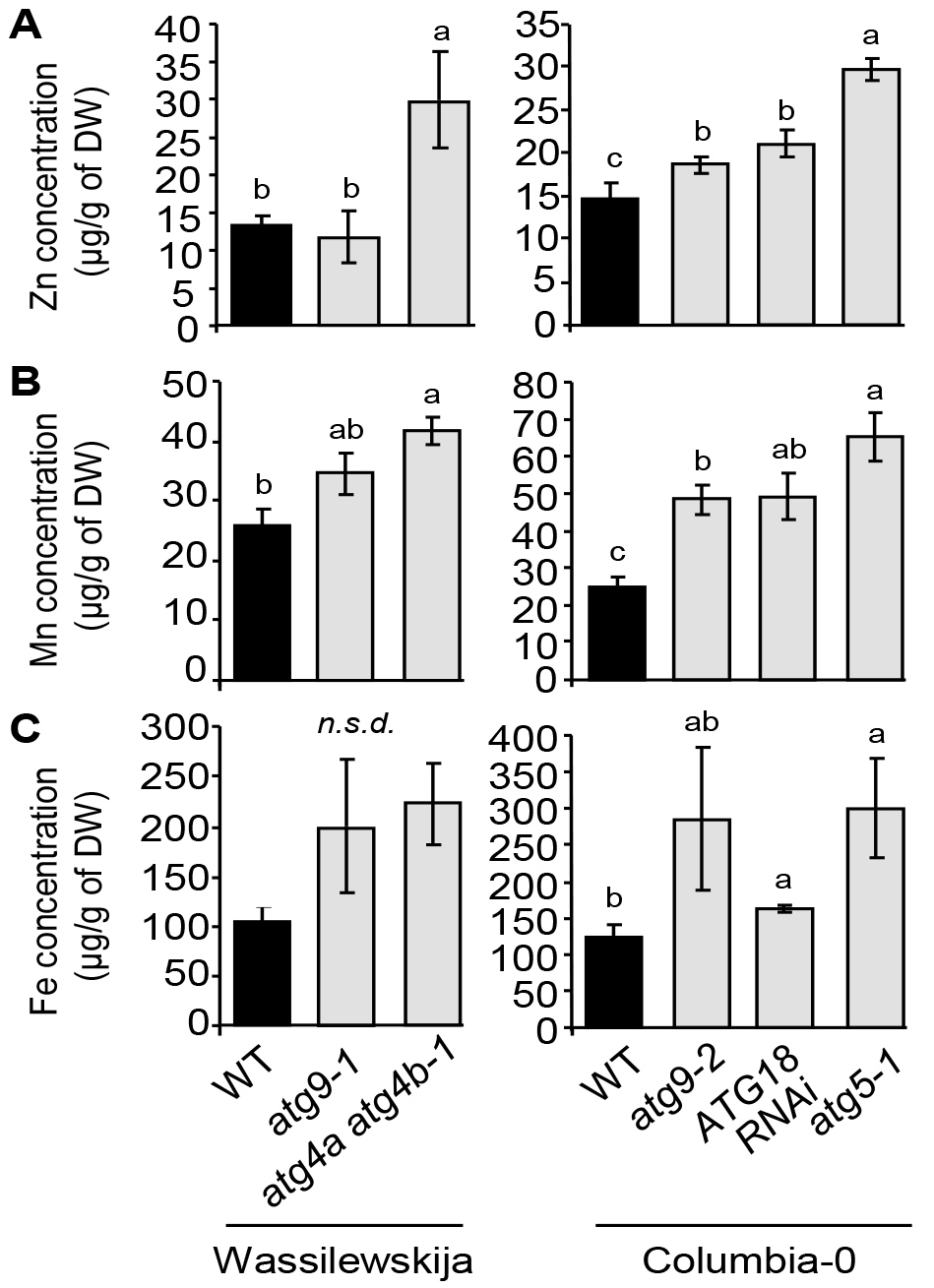
Defects in autophagy lead to elevated Zn, Mn, and Fe concentrations in vegetative tissues of *Arabidopsis thaliana* (experiment 1). Zn (A), Mn (B) and Fe (C) concentrations in dry remains of vegetative tissues of wild-type plants, (Wassilewskija and Columbia-0; black bars) and autophagy deficient plants (gray bars) in the same genetic backgrounds, grown on sand as described in Chardon *et al.* (2010). Results are shown as mean ± SE of four biological repeats. Different letters indicate significant differences between genotypes according to a Kruskal-Wallis test (*p < 0.05*, n = 4) followed by a Tukey *post hoc* test; *n.s.d.* indicates no significant difference.

These data suggest that defects in autophagy affect micronutrient metal resorption from vegetative organs. The severity of the resorption defects appears to be related with the level of autophagy deficiency in the different genotypes tested.

### Distinct mechanisms lead to decrease in Fe concentrations in seeds of autophagy deficient plant depending on Fe supply during the reproductive stage

To further analyze the role of autophagy in micronutrient resorption from vegetative organs to seeds, we chose to focus on the *atg5-1* mutant, which exhibited robust increases in metal concentrations in its dry remains (**Fig. 1**), and on Fe, a critical micronutrient that is often not sufficiently available in seeds of crop plants for human nutrition (Velu *et al.*, 2014). Because autophagy operates a negative feedback loop modulating salicylic acid (SA) signaling to negatively regulate senescence, autophagy deficient plants exhibit a premature senescence phenotype which may interfere with nutrient resorption (Yoshimoto *et al.*, 2009; Guiboileau *et al.*, 2012). Hence, we also included in this experiment (experiment 2) the stay-green mutant *sid2*, which is defective in SA biosynthesis, and *atg5sid2* double mutant, in which autophagy is defective while senescence is delayed (Wildemuth *et al.*, 2001; Yoshimoto *et al.*, 2009).

Seed filling may depend on both remobilization from senescent organs and *de novo* uptake by roots during seed formation. In order to discriminate the two pathways, plants were grown in parallel under two different conditions: some were supplied with sufficient Fe during all their life cycle (Fe sufficient supply), while other were deprived of Fe from the onset of flowering on (Fe deprivation).

Before investigating the contribution of autophagy to Fe remobilization from different organs and subsequent Fe loading in seeds during the reproductive stage, Fe concentrations were first measured in roots and rosette leaves during the vegetative growth of wild-type and *atg5-1* mutant plants (**Supplemental Fig. S1**, experiment 2). No significant difference in Fe concentration was observed between genotypes indicating that Fe uptake and transfer to leaves was not affected in *atg5-1* at this stage (**Supplemental Fig. S1**). Besides, a 3.2 ± 0.6-fold increase in the level of FERRITINS (n = 4; p < 0.05 according to a Mann-Whitney test; **Supplemental Fig. S2**), which are involved in Fe storage in plastids (Briat *et al.*, 2010), was observed at the vegetative stage in *atg5-1* mutant leaves compared to those of wild-type plant. This result suggests that Fe subcellular distribution is modified in *atg5-1*. Fe concentrations were then measured in leaves, stems including empty siliques, and seeds of *atg5-1, atg5sid2, sid2* mutants, and wild-type plants at the end of their life cycle. No differences in Fe concentration were observed between *sid2* and wild-type plants, irrespective of the Fe nutrition regime. However, under Fe sufficient nutrition, Fe concentrations in the *atg5-1* mutant were significantly higher than that of wild-type in leaves (1.8 fold) and in stems (3.4 fold) (**Fig. 2**), in agreement with the results presented in **Figure 1**. The increase in Fe concentration in *atg5-1* vegetative organs was associated with a trend towards decreased seed Fe concentration (**Fig. 2**). Similar results were observed in experiment 3 comparing wild-type and *atg5-1* mutant plants under higher light conditions (**Supplemental Fig. S3**), confirming the robustness of this observation. In experiment 3, metal concentrations were also measured in roots and rosette leaves at the beginning of seed filling (**Supplemental Fig. S4**). Under Fe sufficient conditions, higher Fe concentrations were observed in rosette leaves of *atg5-1* mutant than in those of wild-type plants already at this early stage, whereas no difference was detected at the root level (**Supplemental Fig. S4**). These data suggest that, at least initially, Fe is remobilized from leaves rather than from roots. Both the increase in Fe concentration in vegetative organs and the decrease in seeds observed in *atg5-1* tended to reverse to a large extent in *atg5sid2* (**Fig. 2**, experiment 2), suggesting that these effects are in part due to early leaf senescence.

**Figure 2.**
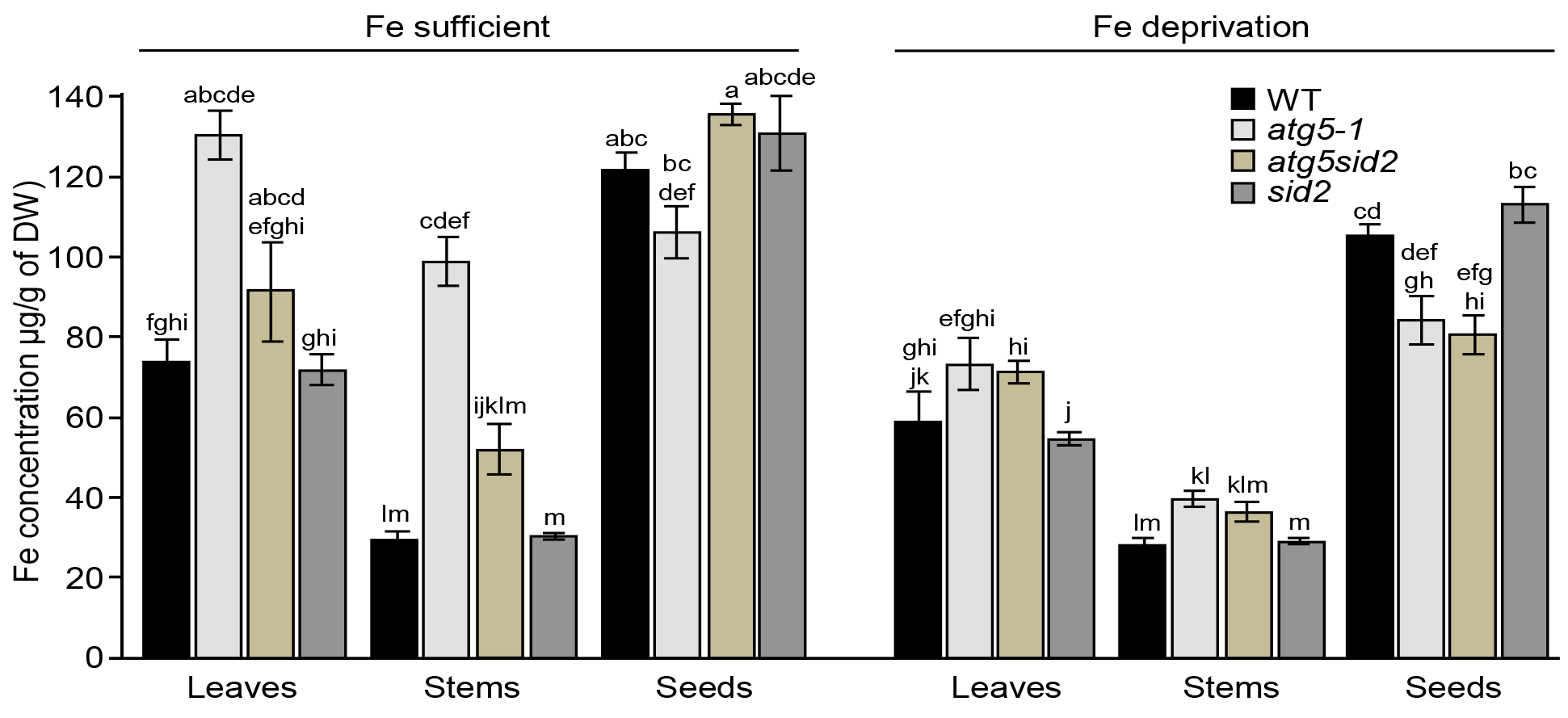
Distinct mechanisms lead to decrease in Fe concentration in seeds of autophagy deficient plant depending on Fe supply during the reproductive stage in *Arabidopsis thaliana* (experiment 2). Fe concentrations in leaves, stems (including empty siliques) and seeds of wild-type (WT, black bars), *atg5-1* mutant (light grey bars), *atg5sid2* double mutant (brown bars) and *sid2* (dark gray bars) mutant plants grown on sand/perlite (1/1) substrate watered with modified Hoagland medium. Iron was supplied (Fe sufficient) or not (Fe deprivation) during the reproductive stage. Results are shown as mean ± SE of five biological repeats. Different letters indicate significant differences according to a Kruskal-Wallis test (*p < 0.05*, n = 5) followed by a Tukey *post hoc* test.

Under Fe deprivation, Fe concentrations were lower in leaves and seeds irrespective of the genotype, as expected (**Fig. 2**). In contrast with the Fe sufficient nutrition, Fe concentrations in vegetative organs of *atg5-1* mutant displayed only a moderate and non-significant increase (1.2 fold) with respect to wild-type, while Fe concentration was significantly decreased by 20 % in seeds (**Fig. 2**). These effects were not reversed in *atg5sid2* mutant, suggesting that they are due to impaired autophagy rather than early senescence.

Taken together, these data indicate that the *atg5-1* mutant is affected in Fe resorption from vegetative organs to seeds by distinct mechanisms under sufficient Fe supply or Fe deprivation during the reproductive period.

### Defective autophagy leads to a strong decrease in the pool of Fe allocated to seeds

To evaluate the impact of autophagy on the redistribution of Fe pools between vegetative organs and seeds, we calculated Fe mass distribution among organs. For this purpose, we first determined the total biomass distribution as well as the harvest index, which is defined as the ratio of seed biomass relative to the whole aerial biomass of the plant after completion of the life cycle. As previously reported (Guiboileau *et al.*, 2012), loss of *ATG5* function led to a strong reduction of total dry biomass (2.8 fold; **Supplemental Fig. S5**) and an even more severe defect in seed production causing a decrease in the harvest index (**Supplemental Fig. S6**). These effects were not significantly reversed in *atg5sid2* mutant, and were also observed when plants were deprived of Fe during the reproductive phase (**Supplemental Fig. S5 and S6**).

Taking biomass and Fe concentrations into account, Fe content [Fe _content_ (μg) = Fe concentration (μg.mg^-1^ of dry weight) x Organ _biomass_ (mg of dry weight)] was determined for each organ (**Fig. 3A**). First, we observed a lower Fe content under Fe deprivation in all genotypes, indicating that *de novo* Fe uptake during seed formation contributes to total Fe content independently of autophagy. (**Fig. 3A**). Although *atg5-1* mutant displayed significantly higher Fe concentrations than wild-type in vegetative organs (**Fig. 2**), its total Fe content was strongly decreased compared to wild-type due to its much lower biomass (**Fig. 3A; Supplemental Fig. S5**). Disrupting *sid2* gene did not restore Fe accumulation in *atg5-1*, indicating that this defect is not due to the premature senescence of *atg5-1* mutant.

**Figure 3.**
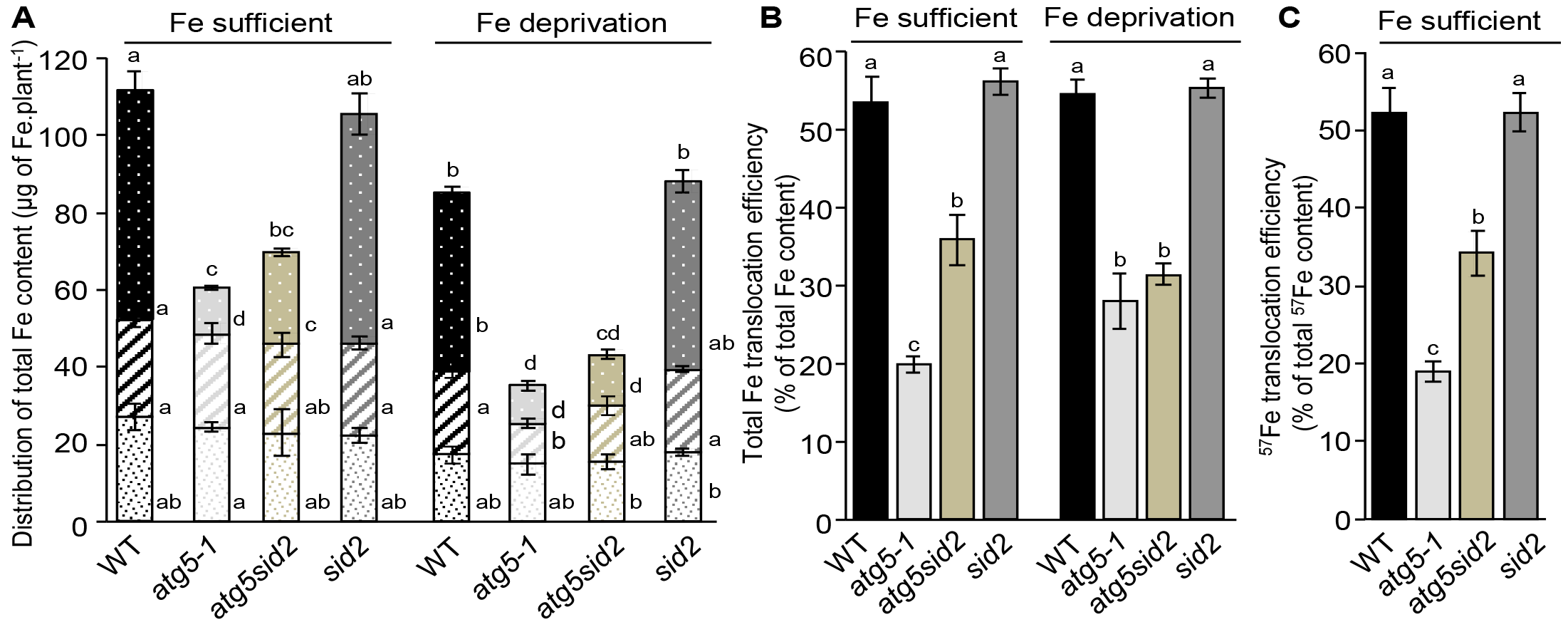
Total Fe content, Fe distribution, and Fe translocation to seeds are dramatically affected in *atg5-1* and *atg5sid2* mutants of *Arabidopsis thaliana* (experiment 2). A, Distribution of Fe between leaves (closed squares), stems including empty siliques (diagonal stripes), and seeds (white dots) of plants are represented for the wild-type (WT, black bars), the *atg5-1* (light gray bars), the *atg5sid2* (brown bars), and the *sid2* (dark gray bars) mutant plants. B, total Fe translocation efficiency calculated as the ratio between the Fe content in seeds and the Fe content in the whole plant (Fe _seed_ / (Fe _leaf_ + Fe _stem_ + Fe _seed_). C, ^57^Fe translocation efficiency (remobilized Fe translocation efficiency) calculated as ^57^Fe_seed_ / (^57^Fe leaf + ^57^Fe _stem_ + ^57^Fe _seed_) after ^57^Fe pulse labeling at the vegetative stage. Plants were grown on sand/perlite (1/1) substrate watered with modified Hoagland medium. Fe was supplied (Fe sufficient) or not (Fe deprivation) during the reproductive stage. Results are shown as mean ± SE of five biological repeats. Different letters indicate significant differences between genotypes and conditions according to a Kruskal-Wallis test (*p < 0.05*, n = 5) followed by a Tukey *post hoc* test.

Compared to that of wild-type, lower Fe contents were observed in vegetative organs of *atg5-1* (-35%) and *atg5sid2* (-24%) mutants specifically under Fe deprivation but not under Fe sufficient nutrition (**Fig. 3A**). This result indicates that Fe supply during the reproductive stage provides vegetative organs of autophagy deficient plants with Fe even though they undergo premature leaf senescence. This suggests that Fe is accumulated in vegetative tissues during the early steps of the reproductive stage.

Iron content of seeds from plants growing under Fe deprivation was dramatically decreased by -80% and -70% in *atg5-1* and *atg5sid2* mutants compared to that of wild-type, respectively (**Fig. 3A**). Iron content is therefore much more severely affected by the lack of autophagy in seeds than in other organs due to a cumulative effect of decreased seed production and decreased seed Fe concentration. Fe supply during the reproductive stage partially restored Fe content of *atg5sid2* seeds but had no effect on the Fe content of *atg5-1* seeds. This result indicates that *de novo* Fe uptake during seed formation can provide seeds with Fe in autophagy deficient plants, but only when premature senescence is prevented. This suggests that such contribution takes place during the late steps of the reproductive stage, when *atg5-1* but not *atg5sid2* displays leaf senescence symptoms.

### Impaired autophagy affects translocation of Fe remobilized from vegetative organs to seeds

We then analyzed the impact of the absence of autophagy on the efficiency of total Fe translocation to seeds (Fe content _seeds_ / Fe content _(leaves + stems + seeds)_), under Fe sufficient nutrition and Fe deprivation (**Fig. 3B**). The efficiency of total Fe translocation to seeds was indistinguishable between wild-type or *sid2* mutant plants representing 50-55% of the total Fe content of the plant, irrespective of the Fe nutrition regime. In contrast, the efficiency of total Fe translocation was strongly decreased in *atg5-1* and *atg5sid2* mutants. Under Fe sufficient nutrition, Fe translocation efficiency was decreased by 63% in *atg5-1* mutant and only 35% in *atg5sid2* mutant confirming that both autophagy and premature senescence limit Fe translocation to seeds under these conditions. Under Fe deprivation, total Fe translocation efficiency was decreased to a similar extent (42-48%) in *atg5-1* and *atg5sid2* mutants, indicating that premature senescence has no impact on total Fe translocation when Fe is not provided during the reproductive phase.

In the experiment 3 performed under higher light conditions, similar total Fe translocation efficiencies of 50-55% were measured in wild-type (**Supplemental Fig. S7**). However, in *atg5-1* mutant, the total Fe translocation efficiency was further decreased under Fe deprivation. We hypothesize that light stress may have modified the timing of leaf senescence or amplified the effect of Fe deficiency.

Using Fe deprivation to estimate the relative contribution of root uptake and remobilization from leaves may be biased by the activation of compensatory mechanisms under Fe deficiency (Pottier *et al.*, 2014). Then, to monitor fluxes specifically between senescing organs and seeds independently of uptake from soil, we performed a ^57^Fe labeling pulse-chase experiment. A pulse of ^57^Fe was provided in the nutritive medium of young rosette plants. Pots and substrate were then thoroughly washed to avoid further ^57^Fe uptake during the chase period. Plants were subsequently watered with nutritive solution containing non-labeled Fe until the end of the life cycle (Fe sufficient nutrition). At the end of the life cycle, the relative abundance of ^57^Fe was determined in each organ by ICP-MS. From these data, we first calculated ^57^FeRSA values which reflect the dilution of ^57^Fe in leaves, stems, and seeds for each genotype (**Supplemental Table S1**). Leaves displayed the highest ^57^FeRSA values (**Supplemental Table S1**), which is consistent with the fact that pulse labeling was performed at the rosette stage when other organs were not present yet. Conversely, stems showed consistently the lowest ^57^FeRSA, suggesting that both Fe remobilization from rosette leaves and *de novo* Fe uptake from soil contributed to stem total Fe (**Supplemental Table S1**). Finally seeds displayed an intermediate ^57^FeRSA level which could reflect mixed contribution of remobilization from leaves and stems, and possibly *de novo* Fe uptake during the reproductive stage. The lower ^57^FeRSA in *sid2* and *atg5sid2* seeds may be due to delayed senescence allowing prolonged uptake of ^56^Fe and further dilution of the label. To obtain further information on the contribution of these different sources of Fe to seed Fe, we calculated the ^57^FeRSA _seeds_ / ^57^FeRSA _(seeds + stems + leaves)_ ratios (**Table 1**), which compares ^57^Fe dilution between the seeds and the whole aerial part of the plant (Gallais *et al.*, 2006). A 57FeRSA _seeds_ / ^57^FeRSA _(seeds + stems + leaves)_ ratio lower than 1 indicates that the ^56^Fe absorbed *de novo* during the reproductive stage contributes mainly to seed Fe content, while a ratio higher than 1 indicates that the ^56^Fe absorbed during the reproductive stage contributes mainly to the Fe content of vegetative tissues. Here, ^57^FeRSA _seeds_ / ^57^FeRSA _(seeds + stems + leaves)_ ratios were almost equal to 1 indicating that ^57^Fe dilution is similar in seeds and in the whole plant, and no significant differences were observed between genotypes (**Table 1**). Such result shows that ^56^Fe taken up during the reproductive stage contributes to a similar extent to Fe content of seeds and to that of the leaves and stems. Two scenarios may account for this value, either (1) direct ^56^Fe fluxes to the vegetative organs and to the seeds are perfectly balanced and lead to similar dilutions or (2) ^56^Fe is first loaded in vegetative organs, which subsequently provide Fe with a predetermined RSA to the seeds. Finally, combining ^57^Fe/^56^Fe ratio with Fe concentration and dry weight, we calculated the ^57^Fe content _seed_ / ^57^Fe content _(leaves + stems +_ seeds) ratio, i.e. the proportion of ^57^Fe translocated from the rosettes and the stems to the seeds. Compared to the apparent Fe translocation ratio, this ratio takes into account Fe taken up only during the vegetative stage and is thus a good indicator of the Fe remobilization efficiency from vegetative organs to seeds even when Fe is supplied during the reproductive stage (**Fig. 3C**). Values of the ^57^Fe content _seed_ / ^57^Fe content _(leaves + stems + seeds)_ ratio were very similar to those of the Fe content _seed_ / Fe content _(leaves + stems + seeds)_ ratio (**Fig. 3B**) irrespective of the genotype. This result suggests that, under our experimental conditions, the loading of Fe to the seeds is mostly achieved by Fe remobilization from the vegetative organs according to scenario (2), even though Fe is still taken up from the soil during the reproductive stage (**Fig. 3A**). Thus, the intermediate ^57^FeRSA level measured in seeds is only due to the mixed contribution of remobilization from vegetative organs (**Supplemental Table 1**) rather than to dilution by ^56^Fe originating directly from root uptake. This is in apparent contradiction with a value of ^57^FeRSA _seeds_ / ^57^FeRSA _(seeds + stems + leaves)_ of 1 indicating that *de novo* uptake during the reproductive growth contributes similarly to Fe content of vegetative organs and seeds (**Table 1**). This apparent discrepancy may be solved if we consider scenario (2) which suggests an indirect involvement of the *de novo* Fe uptake to seed Fe content though a two-step mechanism, as described in the discussion.

**Table 1.** ^57^FeRSA _seeds_ / ^57^FeRSA _(seeds + stems + leaves)_ ratios in wild-type Columbia-0, *atg5-1*, *atg5sid2* and *sid2* mutant plants which have undergone a ^57^Fe pulse labeling at the vegetative stage. No significant differences in ^57^FeRSA _seeds_ / ^57^FeRSA _(seeds + stems + leaves)_ ratios were observed between genotypes according to Kruskal-Wallis test (*p < 0.05*, n = 5).

| Genotype | $^{57}\text{FeRSA}_{\text{seeds}} / ^{57}\text{FeRSA}_{(\text{seeds} + \text{stems} + \text{leaves})}$ |
| --- | --- |
| Col-0 | $0.96 \pm 0.02$ |
| *atg5-1* | $0.98 \pm 0.07$ |
| *atg5sid2* | $0.92 \pm 0.04$ |
| *sid2* | $0.88 \pm 0.04$ |

### The lack of autophagy also impacts Zn and Mn translocation to seeds

Before investigating the putative involvement of autophagy in the translocation of Zn and Mn to seeds, their concentrations were measured in roots and young rosette leaves during the vegetative stage, to test whether *atg5-1* affects Mn or Zn uptake and transfer to leaves at this stage (**Supplemental Fig. S1**). For Zn, no significant differences were observed in leaves and roots as was the case for Fe, indicating similar Zn uptake in both genotypes (**Supplemental Fig. S1**). In experiment 3, metal concentrations were also measured in roots and rosette leaves at the beginning of seed filling (**Supplemental Fig. S4**). At this developmental stage, Zn concentration was significantly higher in both roots and leaves of *atg5-1* mutant than in those of wild-type plants (**Supplemental Fig. S4**). Such result suggests therefore that unlike Fe, autophagy dependent Zn remobilization during seed loading originates from both roots and leaves. Manganese concentration in roots was more than 2 times higher in *atg5-1* mutant compared to wild-type plants at the vegetative stage (**Supplemental Fig. S1,** experiment 2). In experiment 3, root Mn concentration measured at the beginning of seed loading was also much higher in *atg5-1* (**Supplemental Fig. S4**). These observations indicate that disruption of *ATG5* gene strongly perturbs Mn uptake and/or Mn remobilization from roots during both the vegetative and the reproductive growth.

Then, concentrations were studied at the end of the plant life cycle in order to determine Mn and Zn translocation efficiencies under Fe sufficient nutrition, following the same approach as for Fe. In wild-type plants, Zn, and Mn translocation efficiencies were 38% and 24% respectively (**Fig. 4**). In *atg5-1* mutants, total translocation efficiencies of these elements were both strongly affected with respect to wild-type. Similar results were observed in experiment 3 performed under higher light conditions (**Supplemental Fig. S7**). Besides, no significant differences were observed between *atg5-1* and *atg5sid2* mutant plants (**Fig. 4**). Thus, a defect in autophagy affects not only Fe translocation but also at various extents translocation of other micronutrients, even in absence of premature senescence. In view of the results obtained at earlier stages (**Supplemental Fig. S1 and S4**), remobilization from roots should be taken into account to fully evaluate the importance of autophagy for Zn and especially for Mn translocation to seeds. The ratios calculated on the basis of leaf and stem Mn and Zn content (**Fig. 4**) likely underestimate the effect of impairing autophagy.

**Figure 4.**
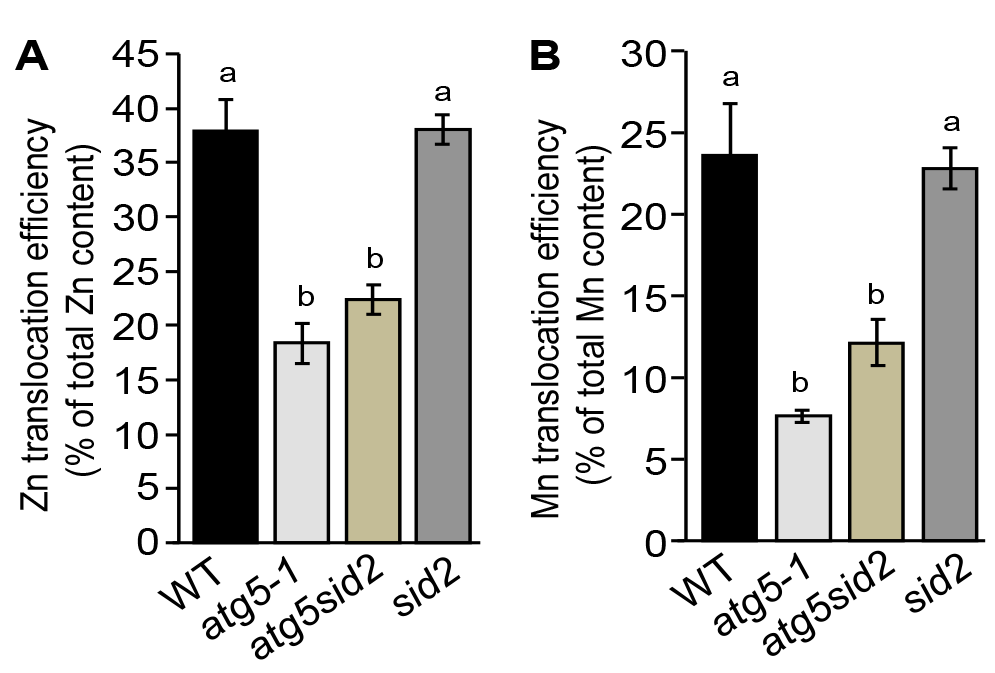
Translocation efficiencies of Zn (A) and Mn (B) are also affected in *atg5-1* and *atg5sid2* mutant plants of *Arabidopsis thaliana* (experiment 2). Translocation efficiencies were calculated for wild-type (WT), *atg5-1*, *atg5sid2*, and *sid2* mutant plants, as the ratio between the micronutrient content in seeds and the micronutrient content in the whole plant. Plants were grown on sand/perlite (1/1) substrate watered with modified Hoagland medium containing Fe during the whole plant cycle. Results are shown as mean ± SE of five biological replicates. Different letters indicate significant differences between genotypes according to a Kruskal-Wallis test (*p < 0.05*, n = 5) followed by a Tukey *post hoc* test.

## Discussion

Because of its poor solubility in most of soils, its essential roles in plant cellular processes, and due to its insufficiency in human diet, many investigations have been undertaken during the last decades to decipher the route of Fe within the plant from its uptake from the rhizosphere to its loading into seeds. Both uptake from soil and remobilization from senescent organs participate to grain loading and thereby contribute to Fe use efficiency (Pottier *et al.*, 2014). To date, little is known about the mechanisms involved in Fe remobilization from senescent source organs during seed formation. A first step of disassembling of organelles, proteins, and various macromolecules is required prior to nutrient reallocation. During the senescence, numerous genes involved in catabolism and degradation mechanisms including genes involved in autophagy are upregulated following a well-established schedule (Breeze *et al.*, 2011). Thus, autophagy may be a limiting step to make nutrients available for further reallocation towards new organs. In the present work, we have investigated the involvement of autophagy in metal micronutrient remobilization from vegetative organs to seeds. Analyzing a range of autophagy mutants, we found that the defect in nutrient remobilization correlated with the severity of the defect in autophagosome formation (**Fig. 1**). Accordingly, the severe defect in autophagy in *atg5-1* mutant leads to a decrease in seed Fe concentration and a drastic reduction in Fe translocation efficiency to seeds (**Figs. 2, 3 and S3**). Interestingly, this defect could be alleviated by providing Fe to the roots only when *atg5-1* premature senescence was prevented (**Fig. 3A**). These results, together with the outcome of the ^57^Fe pulse labeling experiment, suggest that the ^56^Fe taken up by the roots during seed formation contributes indirectly to seed Fe content (**Table 1**; **Figs. 3B and C**). Finally, we found that autophagy plays roles not only in Fe but also in Zn and Mn translocation to seeds, supporting the importance of this mechanism for optimal loading of mineral nutrient to seeds (**Fig. 4**).

### Mineral translocation efficiency to seeds are constants in Arabidopsis

Total iron translocation efficiency of *Arabidopsis thaliana* Columbia-0 wild-type was within the 50-55% range irrespective of light conditions or Fe nutrition regime (**Fig. 3B; Supplemental Fig. S5**). Fe uptake observed during the reproductive stage under Fe sufficient nutrition has therefore no impact on Fe translocation efficiency of wild-type plants (**Fig. 3A and B**). A previous study also found that half of the Fe content of *Arabidopsis* leaves was remobilized during senescence (Himelblau and Amasino, 2001), even though the ecotype, the photoperiod and the growth conditions were different. Thus, Fe translocation efficiency appears to be remarkably stable in *Arabidopsis thaliana*. The other micronutrients studied in this work exhibited lower total translocation efficiencies than Fe (**Fig. 4; Supplemental Fig. S7**). We however found that Zn is better translocated than Mn, which is also in agreement with results obtained by Himelblau and Amasino (2001). In contrast, Maillard *et al.* found large variation in leaf mineral nutrient resorption between different species (Maillard *et al.*, 2015). Pottier *et al.* also found variation in metal resorption from leaves from different poplar genotypes (Pottier et al., 2015). However, remobilization from leaves rather than translocation to seeds was analyzed in these studies.

### Seed iron first transits through vegetative organs

The Fe that is taken up *de novo* during the reproductive stage may be loaded first in young parts of the vegetative organs or directly into seeds. The *atg5-1* mutant, which undergoes premature senescence, displayed similar increase of total Fe content than the other genotypes when Fe is provided during the reproductive stage (**Fig. 3A**). This observation indicates that Fe uptake after the onset of flowering occurs essentially during the early period of the reproductive stage, prior to the senescence of *atg5-1* mutant leaves. Moreover, *atg5-1* mutant accumulated *de novo* absorbed Fe preferentially in vegetative organs while it was translocated more efficiently to seeds when premature senescence was prevented (**Fig. 3A**). If senescence, which eventually leads to cell death, takes place too early, the lapse of time during which nutrients can be translocated between leaves and seeds is shortened, resulting in decreased nutrient translocation efficiency. These results suggest a two-stage process in which Fe taken up by roots is transiently accumulated in vegetative organs prior to its remobilization to the seeds as illustrated in **Figure 5**. This hypothesis also accounts for the similar values observed for total Fe and ^57^Fe remobilization efficiencies as well as for the similar ^57^Fe dilution in seeds and vegetative organs (leaves and stems) reflected in the ^57^FeRSA _seeds_ / ^57^FeRSA _(seeds + stems + leaves)_ ratio almost equal to 1 observed in all genotypes after pulse labeling during the vegetative phase (**Figs. 3B and C; Table 1**). Waters and Grusak (2008) concluded that *de novo* uptake is at least as important for seed filling as remobilization. However, they were using different conditions and did not consider the possibility that minerals have to transit to vegetative organs before being translocated to seeds.

**Figure 5.**
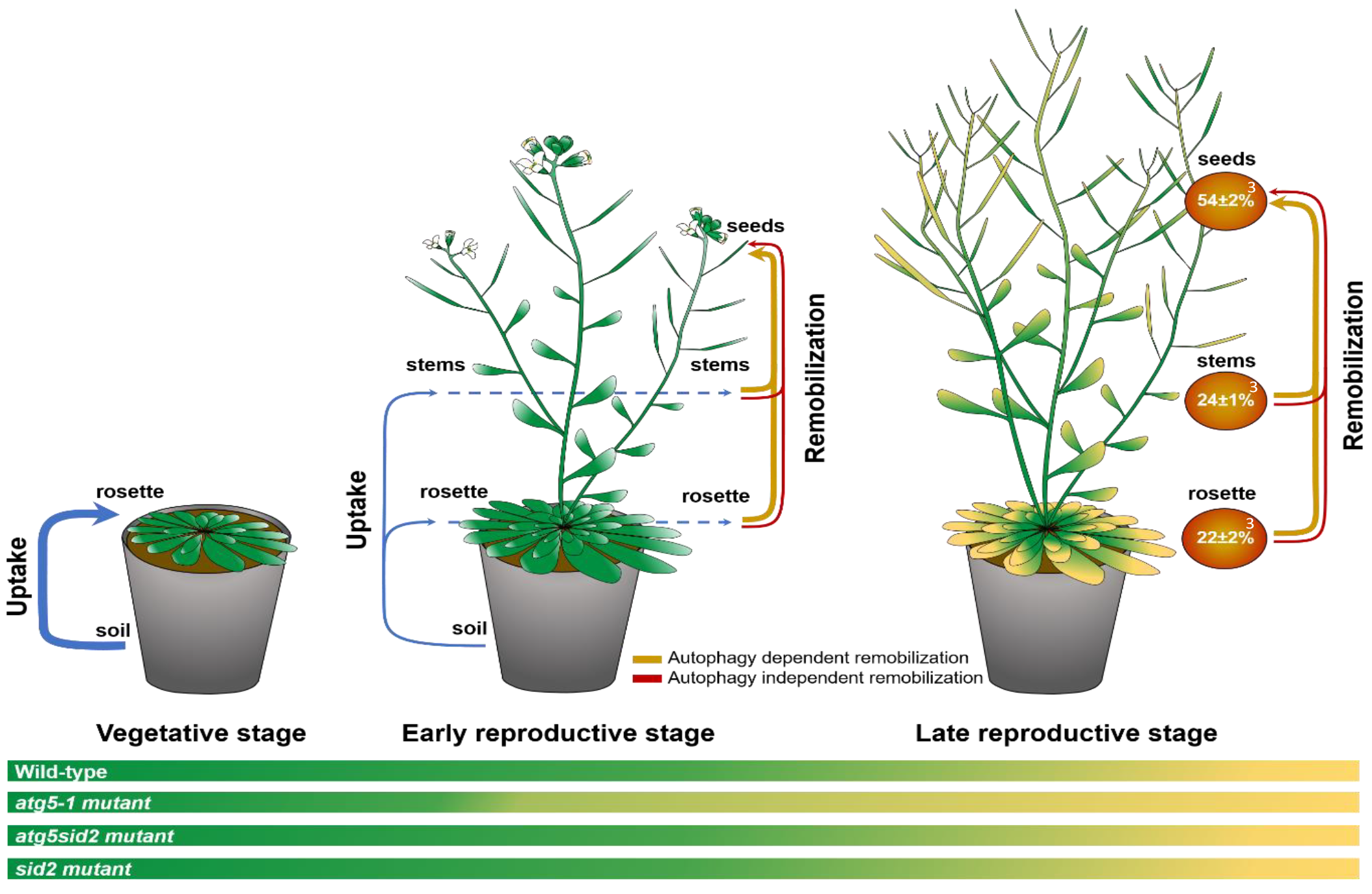
Iron fluxes in wild-type *Arabidopsis thaliana* growing under Fe sufficient condition, in vegetative, early reproductive and late reproductive stages, base of results obtained in wild-type and autophagy deficient plants. Thicknesses of the uptake and remobilization arrows are proportional to flux proportions. Fe distribution (brown circles), are expressed as percentage of total Fe of wild-type plants harvested after completion of the life cycle. Early reproductive stage is defined as the period of premature senescence in *atg5-1* mutant. Late reproductive stage is defined as the period of senescence in the wild-type, when *atg5-1* mutant is already dry. Together, the results obtained indicate that (1) most of the Fe uptake (71±4%^1^) takes place during the vegetative growth, (2) Fe uptake after the onset of flowering (29±5%^2^) occurs mostly during the early reproductive stage, (3) Fe loading into seeds (54±2%^3^) is mainly achieved through remobilization rather than direct contribution from Fe root uptake, (4) in wild-type 35±2%^4^ the total Fe content is loaded to seeds through autophagy dependent remobilization, and 19±2%^5^ is loaded through autophagy independent remobilization. ^1^: total Fe content of plants growing in Fe deprivation condition as a percentage of total iron of plants growing in Fe sufficient condition. ^2^: difference between the total Fe content of plants growing in Fe sufficient condition and the total Fe content of plants growing in Fe deprivation condition as a percentage of total iron of plants growing in Fe sufficient condition. ^3^: Fe content in seeds, stems, and rosette leaves of wild-type plants as percentages of total iron of wild-type plants growing in Fe sufficient condition. ^4^: difference between the percentage of Fe loaded in seeds of wild-type plants and the percentage of Fe loaded in seeds of *atg5sid2* mutant plants. ^5^: difference between the percentage of Fe content in seeds and the percentage of Fe loaded in seeds through autophagy dependent remobilization.

### A possible contribution of root metal micronutrient stores to seed filling

Our work has not addressed the contribution of root Fe stores to total plant Fe and its fate during senescence. The experimental set up that was chosen, growth in a sand perlite mixture, precluded harvesting the whole root system at the end of the plant’s life cycle. Moreover, rigorous analysis of root micronutrient content requires the removal through desorption of cations that are adsorbed on the root but not taken up by living cells. However, at the end of the life cycle the roots are dead, cellular ion content is released and desorption becomes meaningless. Growing plants under hydroponic conditions would allow collection and desorption of the entire root biomass. However, this cultivation system affects nutrient distribution between roots and shoots and is not suitable for seed production (Diaz *et al.*, 2008).

Nevertheless, we collected roots at earlier stages of plant development. Root Fe concentrations were not affected in *atg5-1* mutant at the vegetative stage or at the onset of silique development (**Supplemental Fig. S1 and S4**). This suggests that the defects in Fe distribution observed in the aerial parts of *atg5-1* are not related to root function. Roots are the site of *de novo* Fe uptake as well as a vegetative organ in which Fe may be stored for subsequent remobilization. The results obtained using ^57^Fe labelling argue in favor of a model in which the Fe taken up by roots during the reproductive phase first transits through vegetative organs. However, these results do not exclude a contribution of the Fe stored in roots to seed filling as well as an impact of autophagy defects on remobilization of root Fe pools at later stages. In contrast, differences in Zn concentrations in both rosette leaves and roots at the onset of seed filling suggest that root Zn pools may be reallocated together with leaf Zn pools at the reproductive stage (**Supplemental Fig. S4**). Besides, differences of Mn concentrations were observed in roots at the vegetative stage and at early reproductive stage indicating that autophagy influences its uptake or distribution (Supplemental Fig. S1 and S4). The importance of root Mn pools for Mn redistribution has already been documented in *Brassica napus* subjected to Mn deficiency (Maillard *et al.*, 2015). Regarding nitrogen, remobilization from roots to flowering stems and reproductive organs was shown to have a minor contribution in *A. thaliana* (Diaz *et al.*, 2008). In *Brassica napus*, such remobilization contributes only to 11.1% of the pod nitrogen content (Rossato *et al.*, 2001). Regarding Zn, no decrease of its root content was observed between anthesis and maturity in rice (Jiang *et al.*, 2008), while in wheat, it was decreased by 50% indicating that remobilization of Zn from roots varies according to species (Kutman *et al.*, 2012). In *A. thaliana*, to our knowledge, no evidence for micronutrient remobilization from roots has been reported. Besides, root senescence related to seed loading still has to be investigated (Hammond and White, 2008; Lynch and Brown, 2008). Even though roots were shown to contain higher Fe concentrations than shoots in *A. thaliana* (Cassin *et al.*, 2009; Reyt *et al.*, 2015; Pottier *et al.*, 2015*b*; Eroglu *et al.*, 2016; Li *et al.*, 2016; this study), their contribution to seed filling remains to be demonstrated.

### Harnessing autophagy to improve seed nutrient content

We found that autophagy is a crucial component of micronutrient filling to seeds and thus a strong determinant of seed quality. Although autophagy was shown to be induced during senescence (van der Graaff *et al.*, 2006; Breeze *et al.*, 2011; Htwe *et al.*, 2011), significant amounts of nutrients present in vegetative organs, 30% of N in optimal conditions, 46% of Fe and even larger proportions of other micronutrients, are not reallocated to seeds (**Fig. 4**; Guiboileau *et al.*, 2012). Thus, increasing further autophagy process specifically during seed filling may be a powerful approach to increase the pool of nutrients available for subsequent translocation to seeds. However, it will be also necessary to increase seed storage capacity. In seeds, iron is highly concentrated in vacuoles of the aleurone layer in cereals (Moore *et al.*, 2012) or of the proto-endodermis in several dicot species (Mary *et al.*, 2015; Ibeas *et al.*, 2017). These localizations likely minimize the damage that could be caused by excess free iron. Additional parallel research on the mechanisms of iron storage in seeds is also needed. Moreover, autophagy is a complex mechanism subjected to a subtle multiscale regulation (Yoshimoto, 2012; Masclaux-Daubresse *et al.*, 2017). Further investigations on the regulation of autophagy may thus help improve grain yield by bioengineering strategies or marker assisted breeding.

## Supplementary data

**Supplemental figure S1**. Fe, Mn, and Zn concentrations in roots and rosette leaves measured during the vegetative growth in Col-0 wild-type and *atg5-1* mutant (experiment 2).

**Supplemental figure S2.** FERRITINS are more abundant in vegetative tissues of *atg5-1* mutant than in those of col-0 wild-type (experiment 1).

**Supplemental figure S3**. The increase in Fe concentration in leaves and stems of the *atg5-1* mutant is associated to a significant decrease in seed Fe concentration (experiment 3).

**Supplemental figure S4**. Fe, Mn, and Zn concentrations in roots and rosette leaves measured at the onset of seed loading in Col-0 wild-type and *atg5-1* mutant (experiment 3).

**Supplemental figure S5**. Total dry biomass and dry biomass distribution between organs are dramatically affected in *atg5-1* and *atg5sid2* mutants, even under Fe sufficient condition (experiment 2).

**Supplemental figure S6**. Harvest index is affected in autophagy deficient plants even in absence of premature leaf senescence, independently of the Fe supply condition (experiment 2).

**Supplemental figure S7**. Total seed production and total iron translocation efficiency are dramatically affected in *atg5-1* mutant even under Fe sufficient condition (experiment 3).

**Supplemental figure S8**. Translocation efficiencies of Zn (A) and Mn (B) are affected in *atg5-1* mutant (experiment 3).

**Supplemental table S1.** Relative ^57^Fe-specific allocation (^57^FeRSA) in leaves, stems including empty siliques and seeds, in wild-type Columbia-0, *atg5-1*, *atg5sid2* and *sid2* mutant plants which have undergone a ^57^Fe pulse labeling at the vegetative stage (experiment 2).

## Acknowledgments

The authors are grateful to Cindy Victor and Sara Martins for contribution to harvests and Adrien Tallois and Yann Kourouma for help with elemental analysis. This work was supported by the Centre National de la Recherche Scientifique and the Agence Nationale pour la Recherche grants (ANR-11-BSV6-000 and ANR-16-CE20-0019-02) to S.T. and by the Région Ile-de-France through a grant from the DIM Astréa to M.P. The I2BC and IJPB benefit from the support of the LabEx Saclay Plant Sciences-SPS (ANR-10-LABX-0040-SPS).

**Author contributions:**
S.T., C.M.D., and M.P. designed the experiments. M.P. performed most of the experiments. J.D. developed the analysis of the 56Fe/57Fe ratios and performed isotopic measurements. M.P. and S.T. analyzed the data. M.P. wrote the paper with contributions of all the authors. S.T. supervised and complemented the writing.

## References

Breeze E, Harrison E, McHattie S, et al. 2011. High-resolution temporal profiling of transcripts during Arabidopsis leaf senescence reveals a distinct chronology of processes and regulation. The Plant Cell 23, 873–94.

Briat JF, Duc C, Ravet K, Gaymard F. 2010. Ferritins and iron storage in plants. Biochimica et Biophysica Acta 1800, 806–814.

Cassin G, Mari S, Curie C, Briat JF, Czernic P. 2009. Increased sensitivity to iron deficiency in arabidopsis thaliana overaccumulating nicotianamine. Journal of Experimental Botany 60, 1249–1259.

Chardon F, Barthélémy J, Daniel-Vedele F, Masclaux-Daubresse C. 2010. Natural variation of nitrate uptake and nitrogen use efficiency in Arabidopsis thaliana cultivated with limiting and ample nitrogen supply. Journal of Experimental Botany 61, 2293–2302.

Chung T, Phillips AR, Vierstra RD. 2010. ATG8 lipidation and ATG8-mediated autophagy in Arabidopsis require ATG12 expressed from the differentially controlled ATG12A and ATG12B loci. The Plant Journal 62, 483–493.

Diaz C, Lemaitre T, Christ A, Azzopardi M, Kato Y, Sato F, Morot-Gaudry J-F, Le Dily F, Masclaux-Daubresse C. 2008. Nitrogen recycling and remobilization are differentially controlled by leaf senescence and development stage in Arabidopsis under low nitrogen nutrition. Plant Physiology 147, 1437–1449.

Doelling JH, Walker JM, Friedman EM, Thompson AR, Vierstra RD. 2002. The APG8/12-activating enzyme APG7 is required for proper nutrient recycling and senescence in Arabidopsis thaliana. Journal of Biological Chemistry 277, 33105–33114.

Durrett TP, Gassmann W, Rogers EE. 2007. The FRD3-mediated efflux of citrate into the root vasculature is necessary for efficient iron translocation. Plant Physiology 144, 197–205.

Eroglu S, Meier B, von Wirén N, Peiter E. 2016. The vacuolar manganese transporter MTP8 determines tolerance to iron deficiency-induced chlorosis in Arabidopsis. Plant Physiology 170, 1030–1045.

Fourcroy P, Siso-Terraza P, Sudre D, Saviron M, Reyt G, Gaymard F, Abadia A, Abadia J, Alvarez-Fernandez A, Briat J-F. 2014. Involvement of the ABCG37 transporter in secretion of scopoletin and derivatives by Arabidopsis roots in response to iron deficiency. New Phytologist 201, 155–167.

Gallais A, Coque M, Quilléré I, Prioul J-L, Hirel B. 2006. Modelling postsilking nitrogen fluxes in maize (Zea mays) using 15N-labelling field experiments. New Phytologist 172, 696–707.

van der Graaff E, Schwacke R, Schneider A, Desimone M, Flügge U-I, Kunze R, Flugge U-I, Kunze R. 2006. Transcription analysis of arabidopsis membrane transporters and hormone pathways during developmental and induced leaf senescence. Plant Physiology 141, 776–92.

Guiboileau A, Avila-Ospina L, Yoshimoto K, Soulay F, Azzopardi M, Marmagne A, Lothier J, Masclaux-Daubresse C. 2013. Physiological and metabolic consequences of autophagy deficiency for the management of nitrogen and protein resources in Arabidopsis leaves depending on nitrate availability. New Phytologist 199, 683–94.

Guiboileau A, Yoshimoto K, Soulay F, Bataillé M-P, Avice J-C, Masclaux-Daubresse C. 2012. Autophagy machinery controls nitrogen remobilization at the whole-plant level under both limiting and ample nitrate conditions in Arabidopsis. New Phytologist 194, 732–740.

Hammond J, White P. 2008. Diagnosing phosphorus deficiency in crop plants. The Ecophysiology of Plant-Phosphorus Interactions. Dordrecht, 225–246.

Hanaoka H, Noda T, Shirano Y, Kato T, Hayashi H, Shibata D, Tabata S, Ohsumi Y. 2002. Leaf senescence and starvation-induced chlorosis are accelerated by the disruption of an Arabidopsis autophagy gene. Plant Physiology 129, 1181–1193.

Himelblau E, Amasino RM. 2001. Nutrients mobilized from leaves of Arabidopsis thaliana during leaf senescence. Journal of Plant Physiology 158, 1317–1323.

Htwe NMPS, Yuasa T, Ishibashi Y, Tanigawa H, Okuda M, Zheng S, Iwaya-inoue M. 2011. Leaf senescence of soybean at reproductive stage is associated with induction of autophagy-related genes, GmATG8c, GmATG8i and GmATG4. Plant Production Science 14, 141–147.

Ibeas MA, Grant-Grant S, Navarro N, Perez MF, Roschzttardtz H. 2017. Dynamic subcellular localization of iron during embryo development in brassicaceae seeds. Frontiers in Plant Science 8, 1–8.

Inoue Y, Suzuki T, Hattori M, Yoshimoto K, Ohsumi Y, Moriyasu Y. 2006. AtATG genes, homologs of yeast autophagy genes, are involved in constitutive autophagy in Arabidopsis root tip cells. Plant and Cell Physiology 47, 1641–1652.

Jiang W, Struik PC, Van Keulen H, Zhao M, Jin LN, Stomph TJ. 2008. Does increased zinc uptake enhance grain zinc mass concentration in rice? Annals of Applied Biology 153, 135–147.

Kirisako T, Ichimura Y, Okada H, Kabeya Y, Mizushima N, Yoshimori T, Ohsumi M, Takao T, Noda T, Ohsumi Y. 2000. The reversible modification regulates the membrane-binding state of Apg8/Aut7 essential for autophagy and the cytoplasm to vacuole targeting pathway. The Journal of Cell Biology 151, 263–276.

Kutman UB, Kutman BY, Ceylan Y, Ova EA, Cakmak I. 2012. Contributions of root uptake and remobilization to grain zinc accumulation in wheat depending on post-anthesis zinc availability and nitrogen nutrition. Plant and Soil 361, 177–187.

Lanquar V, Lelièvre FF, Bolte S, et al. 2005. Mobilization of vacuolar iron by AtNRAMP3 and AtNRAMP4 is essential for seed germination on low iron. The EMBO journal 24, 4041–4051.

Lefèvre F, Fourmeau J, Pottier M, Baijot A, Cornet T, Abadía J, Álvarez-Fernández A, Boutry M. 2018. The Nicotiana tabacum ABC transporter NtPDR3 secretes O-methylated coumarins in response to iron deficiency. Journal of Experimental Botany 69, 4419–4431.

Li F, Vierstra RD. 2012. Autophagy: a multifaceted intracellular system for bulk and selective recycling. Trends in Plant Science 17, 526–537.

Li X, Zhang H, Ai Q, Liang G, Yu D. 2016. Two bHLH transcription factors, bHLH34 and bHLH104, regulate iron homeostasis in Arabidopsis thaliana. Plant Physiology 170, 2478 LP–2493.

Liu Y, Bassham DC. 2012. Autophagy: pathways for self-eating in plant cells. Annual Review of Plant Biology 63, 215–237.

Lynch JP, Brown KM. 2008. Root strategies for phosphorus acquisition. The ecophysiology of plant-phosphorus interactions. Dordrecht, 83–116.

Maillard A, Diquélou S, Billard V, Laîné P, Garnica M, Prudent M, Garcia-Mina J-M, Yvin J-C, Ourry A. 2015. Leaf mineral nutrient remobilization during leaf senescence and modulation by nutrient deficiency. Frontiers in Plant Science 6, 317.

Mary V, Schnell Ramos M, Gillet C, et al. 2015. Bypassing iron storage in endodermal vacuoles rescues the iron mobilization defect in the natural resistance associated-macrophage protein3natural resistance associated-macrophage protein4 double mutant. Plant Physiology 169, 748–759.

Masclaux-Daubresse C, Chardon F. 2011. Exploring nitrogen remobilization for seed filling using natural variation in Arabidopsis thaliana. Journal of Experimental Botany 62, 2131–2142.

Masclaux-Daubresse C, Chen Q, Havé M. 2017. Regulation of nutrient recycling via autophagy. Current Opinion in Plant Biology 39, 8–17.

Moore KL, Zhao FJ, Gritsch CS, Tosi P, Hawkesford MJ, McGrath SP, Shewry PR, Grovenor CRM. 2012. Localisation of iron in wheat grain using high resolution secondary ion mass spectrometry. Journal of Cereal Science 55, 183–187.

Morrissey J, Baxter IR, Lee J, Li L, Lahner B, Grotz N, Kaplan J, Salt DE, Guerinot M Lou. 2009. The ferroportin metal efflux proteins function in iron and cobalt homeostasis in Arabidopsis. The Plant Cell 21, 3326–38.

Murgia I, Arosio P, Tarantino D, Soave C. 2012. Biofortification for combating ‘hidden hunger’for iron. Trends in Plant Science 17, 47–55.

Nouet C, Motte P, Hanikenne M. 2011. Chloroplastic and mitochondrial metal homeostasis. Trends in Plant Science 16, 395–404.

Pottier M, Garcia de la Torre VS, Victor C, David LC, Chalot M, Thomine S. 2015a. Genotypic variations in the dynamics of metal concentrations in poplar leaves: A field study with a perspective on phytoremediation. Environmental Pollution 199, 73–82.

Pottier M, Masclaux Daubresse C, Yoshimoto K, Thomine S. 2014. Autophagy as a possible mechanism for micronutrient remobilization from leaves to seeds. Frontiers in Plant Science 5, 11.

Pottier M, Oomen R, Picco C, Giraudat J, Scholz-Starke J, Richaud P, Carpaneto A, Thomine S. 2015b. Identification of mutations allowing Natural Resistance Associated Macrophage Proteins (NRAMP) to discriminate against cadmium. The Plant Journal 83, 625–37.

Reumann S, Voitsekhovskaja O, Lillo C. 2010. From signal transduction to autophagy of plant cell organelles: lessons from yeast and mammals and plant-specific features. Protoplasma 247, 233–256.

Reyt G, Boudouf S, Boucherez J, Gaymard F, Briat JF. 2015. Iron- and ferritin-dependent reactive oxygen species distribution: Impact on arabidopsis root system architecture. Molecular Plant 8, 439–453.

Robinson NJ, Procter CM, Connolly EL, Guerinot M Lou. 1999. A ferric-chelate reductase for iron uptake from soils. Nature 397, 694–697.

Rossato L, Laine P, Ourry A. 2001. Nitrogen storage and remobilization in Brassica napus L. during the growth cycle: nitrogen fluxes within the plant and changes in soluble protein patterns. Journal of Experimental Botany 52, 1655–63.

Santi S, Schmidt W. 2009. Dissecting iron deficiency-induced proton extrusion in Arabidopsis roots. New Phytologist 183, 1072–1084.

Schuler M, Rellán-Álvarez R, Fink-Straube C, Abadía J, Bauer P. 2012. Nicotianamine functions in the phloem-based transport of iron to sink organs, in pollen development and pollen tube growth in Arabidopsis. The Plant Cell 24, 2380–400.

Shi R, Weber G, Köster J, Reza-hajirezaei M, Zou C, Zhang F, Von Wirén N. 2012. Senescence-induced iron mobilization in source leaves of barley (Hordeum vulgare) plants. New Phytologist 195, 372–383.

Shingles R, North M, McCarty RE. 2002. Ferrous ion transport across chloroplast inner envelope membranes. Plant Physiology 128, 1022–1030.

Stacey MG, Patel A, McClain WE, Mathieu M, Remley M, Rogers EE, Gassmann W, Blevins DG, Stacey G. 2008. The Arabidopsis AtOPT3 protein functions in metal homeostasis and movement of iron to developing seeds. Plant Physiology 146, 589–601.

Thompson AR, Doelling JH, Suttangkakul A, Vierstra RD. 2005. Autophagic nutrient recycling in Arabidopsis directed by the ATG8 and ATG12 conjugation pathways. Plant Physiology 138, 2097–2110.

Velu G, Ortiz-Monasterio I, Cakmak I, Hao Y, Singh RP. 2014. Biofortification strategies to increase grain zinc and iron concentrations in wheat. Journal of Cereal Science 59, 365–372.

Vert G, Grotz N, Dedaldechamp F, Gaymard FF, Guerinot M Lou, Briat J-FJ-F, Curie C. 2002. IRT1, an Arabidopsis transporter essential for iron uptake from the soil and for plant growth. The Plant Cell 14, 1223–33.

Waters BM, Grusak MA. 2008. Whole-plant mineral partitioning throughout the life cycle in Arabidopsis thaliana ecotypes Columbia, Landsberg erecta, Cape Verde Islands, and the mutant line ysl1ysl3. New Phytologist 177, 389–405.

WHO. 2016. Iron Deficiency Anaemia.

Wildemuth MC, Dewdney J, Wu G, Ausubel FM. 2001. Isochorismate synthase is required to synthetize salicylic acid for plant defense. Nature 414, 562–565.

Xiong Y, Contento AL, Bassham DC. 2005. AtATG18a is required for the formation of autophagosomes during nutrient stress and senescence in Arabidopsis thaliana. The Plant Journal 42, 535–546.

Yoshimoto K. 2012. Beginning to understand autophagy, an intracellular self-degradation system in plants. Plant and Cell Physiology 53, 1355–1365.

Yoshimoto K, Hanaoka H, Sato S, Kato T, Tabata S, Noda T, Ohsumi Y. 2004. Processing of ATG8s, ubiquitin-like proteins, and their deconjugation by ATG4s are essential for plant autophagy. The Plant Cell 16, 2967–2983.

Yoshimoto K, Jikumaru Y, Kamiya Y, Kusano M, Consonni C, Panstruga R, Ohsumi Y, Shirasu K. 2009. Autophagy negatively regulates cell death by controlling NPR1-dependent salicylic acid signaling during senescence and the innate immune response in Arabidopsis. The Plant Cell 21, 2914–27.

